# Predictive entrainment of natural speech through two fronto-motor top-down channels

**DOI:** 10.1101/280032

**Authors:** Hyojin Park, Gregor Thut, Joachim Gross

## Abstract

Natural communication between interlocutors is enabled by the ability to predict upcoming speech in a given context. Previously we showed that these predictions rely on a fronto-motor top-down control of low-frequency oscillations in auditory-temporal brain areas that track intelligible speech. However, a comprehensive spatio-temporal characterisation of this effect is still missing. Here, we applied transfer entropy to source-localised MEG data during continuous speech perception. First, at low frequencies (1-4 Hz, brain delta phase to speech delta phase), predictive effects start in left fronto-motor regions and progress to right temporal regions. Second, at higher frequencies (14-18 Hz, brain beta power to speech delta phase), predictive patterns show a transition from left inferior frontal gyrus via left precentral gyrus to left primary auditory areas. Our results suggest a progression of prediction processes from higher-order to early sensory areas in at least two different frequency channels.

## Introduction

Natural communication between interlocutors may seem effortless, however it relies on a series of complex computational tasks that have to be performed in the human brain in real-time and often in the presence of noise and other interferences. This high performance can only be achieved with the help of very effective prediction mechanisms (Levinson, 2016; Norris, McQueen, & Cutler, 2016). As auditory speech signals enter the sensory auditory system and are complemented by visual signals and cues, the human brain generates and constantly updates predictions about the timing and content of upcoming speech (Friston and Frith, 2015; Pickering and Garrod, 2007, 2013). In this context of natural conversation, we can model human brains as dynamic systems that are coupled through sensory information and operate according to active inference principles (Friston and Frith, 2015). In this framework, the brain relies on internal models to generate predictions about itself and others and updates the internal model to minimize prediction errors.

The temporal structure of this predictive coding mechanism can be mediated by cortical oscillations and previous studies have shown the computational role of cortical oscillations in speech processing as critical elements for parsing and segmentation of connected speech not only for auditory speech (Arnal and Giraud, 2012; Ding, Melloni, Zhang, Tian, & Poeppel, 2016; Giraud and Poeppel, 2012) but also for visual speech (Giordano et al., 2017; Park, Kayser, Thut, & Gross, 2016; Zion Golumbic, Cogan, Schroeder, & Poeppel, 2013). In addition, cortical oscillations track hierarchical components of speech rhythms and cortical oscillations themselves are hierarchically nested for speech tracking (Gross, Hoogenboom, et al., 2013). Importantly, these findings are evident only for intelligible speech processing where top-down modulation by prediction is possible. In our previous study, we found that low-frequency rhythms in the left frontal and motor cortices carry top-down signals to sensory areas, particularly to left auditory cortex, and this top-down signal was correlated with entrainment to speech (Park, Ince, Schyns, Thut, & Gross, 2015).

While previous studies have provided convincing evidence that low-frequency brain rhythms are involved in mediating top-down predictions, several important questions are still unresolved. First, what is the spatio-temporal structure of these prediction processes, or put differently, when and where in the brain is neural activity predictive of upcoming speech in an intelligibility-dependant manner? Second, what is the relationship between these low-frequency rhythms and higher frequency rhythms that have been implicated in prediction (Morillon and Baillet, 2017)? Third, how do predictive processes (preceding speech) interact with reactive processes (following speech)?

In order to address these questions, we used causal connectivity analysis - transfer entropy (TE) - to identify directed coupling between brain rhythms and speech rhythms for a range of positive delays (brain activity following speech) and negative delays (brain activity preceding speech). For speech signal, we analysed low-frequency phase information which is a dominant spectral component in speech (Chandrasekaran, Trubanova, Stillittano, Caplier, & Ghazanfar, 2009). For brain signal, we analysed both low-frequency phase information as well as high frequency beta power. We hypothesized that beta rhythm in the brain, particularly in higher order areas, are involved in the prediction of forthcoming speech as suggested by the role of beta oscillations in top-down predictive mechanism (Arnal and Giraud, 2012; Bastos et al., 2012; Fontolan, Morillon, Liegeois-Chauvel, & Giraud, 2014). A recent EEG study that used time-compressed speech reported beta oscillations reflecting an endogenous top-down channel which gradually builds up contextual information across time (Pefkou, Arnal, Fontolan, & Giraud, 2017). Particularly, beta oscillations in motor system were shown to be associated with precise temporal anticipation of forthcoming auditory inputs (Morillon and Baillet, 2017). We also hypothesized that this top-down ‘predictive speech coding’ mechanism by beta oscillations (which should be represented at negative delays between brain activity and speech) recurrently interacts with low-frequency ‘speech entrainment’ (represented at positive delays) where better prediction leads to stronger entrainment.

## Materials and Methods

### Participants and Experiment

Twenty-two volunteers participated in the study (11 females; age range 19–44 years, mean age ± SD: 27.2 ± 8.0 years). None of the participants had a history of psychological, neurological, or developmental disorders. They all had normal or corrected-to-normal vision and were right-handed. Written informed consent was obtained from all participants prior to the experiment and all participants received monetary compensation for their participation. The study was approved by the local ethics committee (FIMS00733; University of Glasgow, Faculty of Information and Mathematical Sciences) and conducted in accordance with the Declaration of Helsinki.

Participants were instructed to listen to a recording of a 7-min-long story (“Pie-man,” told by Jim O‘Grady at “The Moth” storytelling event, New York). The stimulus was presented binaurally via a sound pressure transducer through two 5-meter-long plastic tubes terminating in plastic insert earpieces. Stimulus presentation was controlled via Psychtoolbox (Brainard, 1997) in MATLAB (MathWorks, Natick, MA). The experiment consisted of two conditions: standard presentation of story (intelligible speech) and backward played presentation of story (unintelligible speech). Experimental conditions were presented in randomised order across participants.

### Data acquisition, Preprocessing and Source localisation

Data recordings were acquired with a 248-magnetometers whole-head MEG system (MAGNES 3600 WH, 4-D Neuroimaging) in a magnetically shielded room. Data were sampled at 1017 Hz and resampled at 250 Hz, denoised with information from the reference sensors, and detrended. The analysis was performed using the FieldTrip toolbox (Oostenveld, Fries, Maris, & Schoffelen, 2011) (http://fieldtrip.fcdonders.nl) and in-house MATLAB scripts according to the guidelines (Gross, Baillet, et al., 2013).

Structural T1-weighted magnetic resonance images (MRI) of each participant were obtained at 3T Siemens Trio Tim Scanner (Siemens, Erlangen, Germany) and co-registered to the MEG coordinate system using a semi-automatic procedure. Anatomical landmarks (nasion, left and right pre-auricular points) were manually identified in the individual‘s MRI. Both coordinate systems were initially aligned based on these three points. Numerical optimisation by using the iterative closest point (ICP) algorithm (Besl and McKay, 1992) was applied.

Individual MRIs were segmented to grey matter, white matter, and cerebrospinal fluid to create individual head models. Leadfield computation was based on a single shell volume conductor model (Nolte, 2003) using a 10-mm grid defined on the standard template brain (Montreal Neurological Institute; MNI). The template grid was transformed into individual head space by linear spatial transformation. Cross-spectral density matrices were computed using Fast Fourier Transform on 1-s segments of data after applying Hanning window.

Frequency-specific spatial filters were computed for delta (1-4 Hz) and beta (14-18 Hz) bands at each voxel. We computed the covariance matrix over the full broad-band 7-minute data to compute LCMV filters for each voxel using 7% regularization. These time series were then subjected to band-pass filtering (4th order Butterworth filter, forward and reverse). Dominant dipole orientation was estimated using SVD at each voxel. Bandpass filtered data were projected through the filter to obtain band-limited time-series for each voxel. This computation was performed for each frequency band, and each experimental condition (intelligible and unintelligible). Hilbert transformation was applied to obtain instantaneous phase and power. We also used regions of interest (ROI) maps from the AAL (Automated Anatomical Labeling) atlas (Tzourio-Mazoyer et al., 2002) in order to delineate the temporal characteristics of directed causal relationship over the delays (see below) within the anatomically parcellated regions. We used ROIs labelled Heschl gyrus, inferior frontal gyrus – opercular part, and precentral gyrus.

### Directed causal connectivity analysis by Transfer Entropy (TE)

In this paper, we aimed to investigate two relationships of speech entrainment and predictive speech coding between speech and brain signal. In order to assess the relationships, we used transfer entropy (TE) that quantifies directed statistical dependencies between two signals, i.e., time-lagged predictability. TE is also known as Directed Information (Ince, Schultz, & Panzeri, 2014; Massey, 1990; Schreiber, 2000).

For speech entrainment, we computed TE from speech signal to signal at each brain voxel to quantify to what extent knowledge of speech signal reduces uncertainty in predicting the future of brain signal over and above what could be predicted from knowledge of the past of brain signal alone. For predictive speech coding, we computed TE from brain signal at each voxel to speech signal to quantify to what extent knowledge of brain signal reduces uncertainty in predicting the future of speech signal over and above what could be predicted from knowledge of the past of speech signal alone.

Specifically, we quantized the phase values from the two signals across all time points during stimulus presentation, and then used 4 bins in which each bin was equally occupied. For a specific delay d, we computed TE from speech (*X*) to brain (*Y*) for speech entrainment, and from brain (*X*) to speech (*Y*) for predictive speech coding as follows:

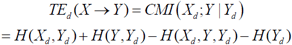

Where CMI is conditional mutual information, H represents entropy. The suffix *d* represents that signal is delayed with respect to the target signal *Y* by *d* milliseconds (i.e. considers that signal *d* milliseconds prior to *Y*). We computed entropy terms from the standard formula:

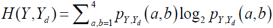

Where the joint distribution *p*_*Y, Yd*_ (*a,b*) is obtained from the multinomial maximum likelihood estimate obtained over time points:

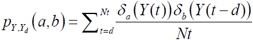

With *δ*_*a*_ (*Y* (*t*)) a Kronecker delta function taking the value 1 if the binned phase value at *Y*(*t*) is quantile a and 0 otherwise.

In the TE computation bias correction was not applied. Bias of mutual information depends on the number of bins and time points that are used in the analysis (Panzeri, Senatore, Montemurro, & Petersen, 2007). In our analysis, we performed statistical contrast between conditions in which the same number of bins and time points was used for each calculation (Ince, Mazzoni, Bartels, Logothetis, & Panzeri, 2012). Bias correction reduces bias but increases the variance of the estimator, so comparisons between calculations with the same bias are better with uncorrected estimates.

The TE calculation was repeated for 25 different delays, from 20 ms to 500 ms with a 20-ms step. These computations were performed for each participant, and both conditions (intelligible, unintelligible). For TE computation to study speech entrainment (TE from speech to brain), we analysed the same frequency (delta; 1-4 Hz) phase information for both speech and brain signals. For TE computation to study predictive speech coding (TE from brain to speech), we analysed 1) the same delta (1-4 Hz) phase information for both brain and speech signals as well as 2) beta (14-18 Hz) power for brain signal and delta (1-4 Hz) phase for speech signal. These computations resulted in TE values for each voxel, each delay, and each condition within each participant and then yielded to statistical comparison between conditions for each delay. For ROI analysis using AAL atlas map, TE values were first averaged across the voxels within the ROI, and then yielded to statistics.

Group statistics was performed using non-parametric randomisation in FieldTrip (Monte Carlo randomisation). Specifically, individual volumetric maps were smoothed with a 10-mm Gaussian kernel and subjected to dependent-samples t-test between conditions (intelligible versus unintelligible). The null distribution was estimated using 500 randomisations and multiple comparison correction was performed using FDR.

## Results

To investigate the bidirectional nature of speech-brain coupling during listening to continuous speech we computed transfer entropy (TE) – an information-theoretic measure of directed causal connectivity - between speech and brain signals at positive and negative delays. This allows a disambiguation of two different effects that contribute to speech-brain coupling – namely those that follow speech (entrainment) and those that precede speech (prediction).

Since speech-brain coupling is strongest for the low frequency delta rhythm (1-4 Hz) we performed our analysis for this frequency band. In the following, we first present results for positive delays where delta-band brain activity follows speech. Second, we show how delta-band brain activity at different negative delays (i.e. preceding speech) in different brain areas predicts (in a statistical sense) the speech signal. Finally, since beta oscillations in motor cortex have been implicated in temporal predictions (e.g. (Arnal and Giraud, 2012; Morillon and Baillet, 2017) we investigate how these higher frequency oscillations in the brain are related to the low-frequency delta rhythm in speech.

### Entrained brain signals following speech (positive delays)

We first examine how low-frequency speech signal entrains the same frequency brain rhythms with positive delays (brain signals following speech signals, Figure 2A). We computed TE from speech envelope to brain signals from different ROIs with delays ranging from 20 ms to 500 ms in steps of 20 ms. Based on our recent study (Park, et al., 2015), we focused on three ROIs from the AAL atlas; primary auditory cortex, inferior frontal gyrus (IFG) and precentral gyrus. We computed TE between intelligible (forward played) and unintelligible (backward played) speech conditions (Figure 2B–D). Whole brain results of the same analysis are shown in Supplementary Figure 1.

**Figure 1:**
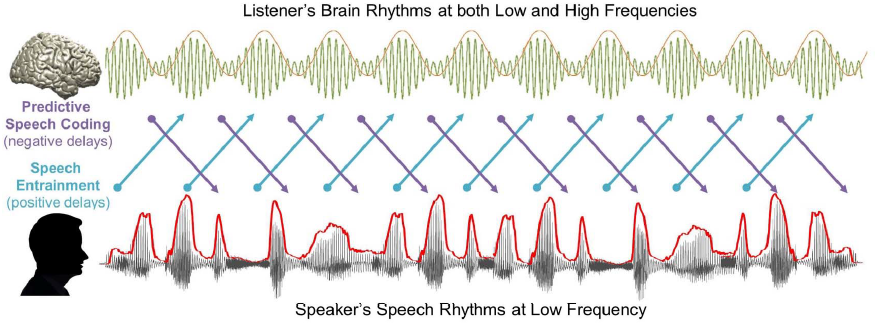
Temporal dynamics of information flow during natural speech perception. In the present study, we analysed transfer entropy (TE) to investigate two mechanisms of information flow during natural speech perception: 1) Predictive speech coding (top-down prediction mechanism investigated by negative delays between speech-brain; purple color) and 2) Speech entrainment (stimulus-driven bottom-up processing investigated by positive delays between speech-brain; cyan color). For speech signal, we used low-frequency delta phase information, and for brain signal, we used both low-frequency delta phase and high-frequency beta power information.

**Figure 2:**
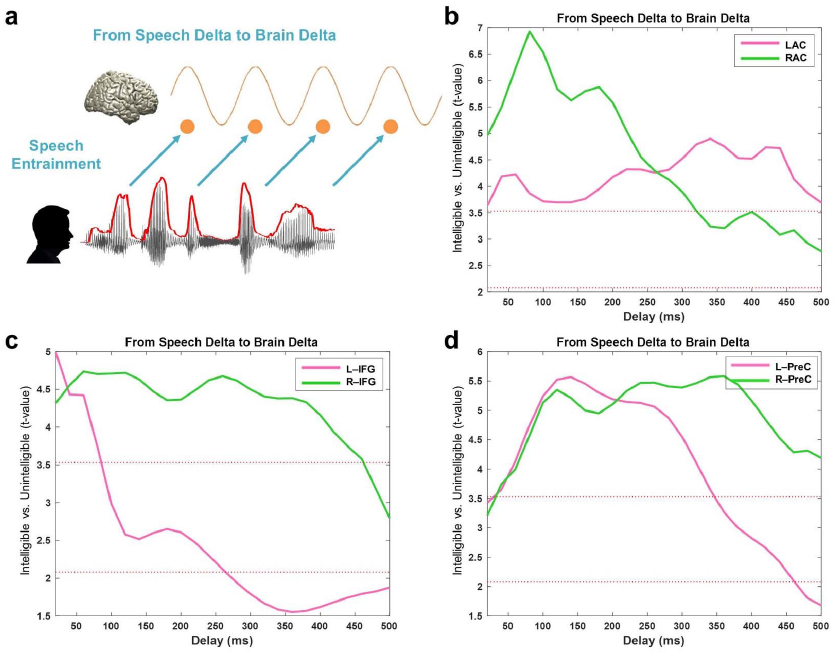
Entrained brain signals following speech. (A) A schematic figure for directed causal analysis: TE from speech delta phase to brain delta phase (positive delays between speech-brain). TE computation was performed for each condition (intelligible-forward played and unintelligible-backward played) at each voxel from 20 ms to 500 ms with a 20-ms step. TE values are averaged within each ROI from the AAL atlas and compared statistically between conditions. T-values are shown in each ROI bilaterally (pink: left hemisphere, green: right hemisphere): (b) primary auditory cortex (Heschl gyrus) (c) Inferior frontal gyrus – opercular part (BA44) (d) precentral gyrus. Statistical significance was shown with two red lines depicting t-values by paired t-test (upper red line: t_21_ = 3.53, p < 0.05, corrected; bottom red line: t_21_ = 2.08, p < 0.05, uncorrected).

Figure 2B shows the statistical difference between intelligible and unintelligible speech across different delays for left (pink) and right (green) auditory cortex. Highest t-values are observed at delays of about 80 ms in right auditory cortex and are stronger than corresponding effects in left auditory cortex. The effect of intelligibility remains significant for delays up to about 300 ms for right auditory cortex and for delays up to 500 ms for left auditory cortex. In IFG, the effect of intelligibility is also most pronounced in the right hemisphere (up to about 450 ms) compared to only early transient effect for the left hemisphere (Figure 2C). Precentral areas in left and right hemisphere show similar sensitivity to intelligibility but are longer lasting in the right hemisphere (Figure 2D).

### Entrained brain signals predicting upcoming speech (negative delays)

Next, we aimed to characterise the spatio-temporal pattern of predictive processes in the brain in the same low-frequency band (1-4 Hz, Figure 3A). Specifically, we computed TE between speech and brain signals over a range of negative delays (−500 ms to −20 ms in steps of 20 ms) to assess where and when brain signals predict significantly stronger upcoming intelligible compared to unintelligible speech. We performed the computation in the whole brain and show maps of statistical difference between intelligible and unintelligible speech condition. Overall, we found the strongest directed effect in left fronto-motor regions ~−220 ms prior to the forthcoming speech when speech is intelligible compared to unintelligible (Figure 3B; paired t-test; upper red line: t_21_ = 3.53, p < 0.05, corrected; bottom red line: t_21_ = 2.08, p < 0.05, uncorrected).

**Figure 3:**
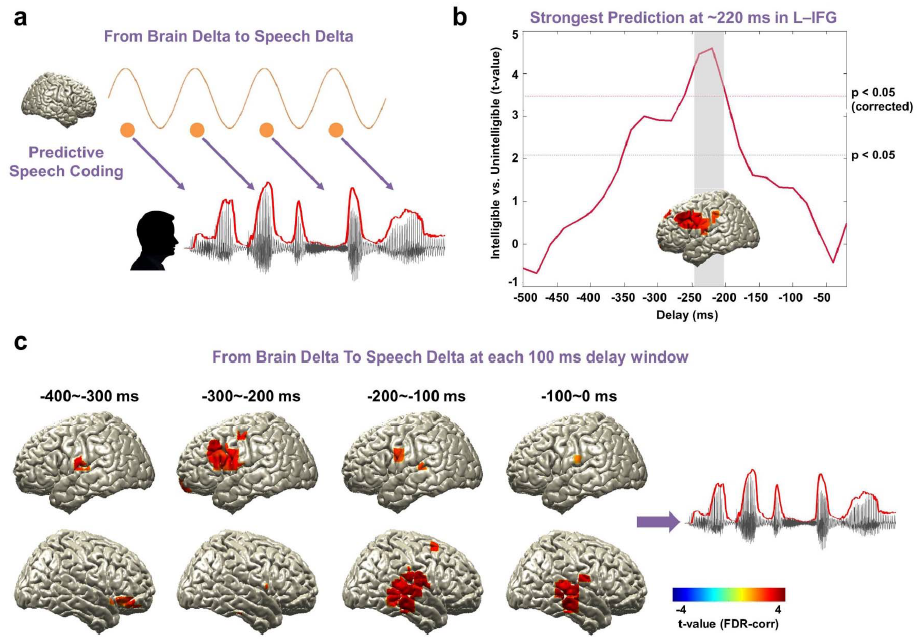
Entrained brain signals predicting upcoming speech. (A) A schematic figure for directed causal analysis: TE from brain delta phase to speech delta phase (negative delays between speech-brain). TE computation was performed for each condition (intelligible-forward played and unintelligible-backward played) at each voxel from 20 ms to 500 ms in 20-ms steps and compared statistically between conditions. (b) The strongest prediction was found at ~220 ms in left IFG (upper red line: t_21_ = 3.53, p < 0.05, corrected; bottom red line: t_21_ = 2.08, p < 0.05, uncorrected). (c) Statistical contrast maps of averaged across 100 ms windows show a sequence of events that start around −300 ms in fronto-motor areas and then move to right auditory-temporal areas at around −200 ms prior to speech (p < 0.05, FDR-corrected).

To further characterise the spatio-temporal evolution of brain signals that predict upcoming speech, we averaged TE-maps in 100 ms windows and computed again the statistical contrast of intelligible compared to unintelligible speech (Figure 3C). This revealed a sequence of events that starts around −300 ms in fronto-motor areas (see also Figure 3B) and then moves to right auditory-temporal areas at around −200 ms prior to speech (Figure 3C; p < 0.05, FDR-corrected). These results indicate that brain activity in fronto-motor areas preceding speech by 300 ms contain information that predicts the forthcoming speech significantly better for intelligible compared to unintelligible speech. The same holds true for right auditory-temporal areas at shorter delays of 200 ms preceding speech. This suggests that predictive speech mechanisms occur first in left inferior fronto-motor areas and later in right auditory-temporal areas.

### Beta rhythms and the prediction of upcoming speech (negative delays)

As we hypothesized based on the literature that beta rhythms are involved in predictive process during speech processing, we examined the causal relationship between beta rhythm in the brain and low-frequency rhythm in the speech signal. We computed TE from beta (14-18 Hz) power at each voxel to the low-frequency delta (1-4 Hz) phase of the speech signal for a range of delays (−500 ms to −20 ms in steps of 20 ms, Figure 4A). The same computation was performed for the intelligible and unintelligible conditions and then statistically compared. We first focus on the temporal dynamics of the three ROIs (primary auditory cortex (Heschl gyrus), inferior frontal gyrus, precentral gyrus).

**Figure 4:**
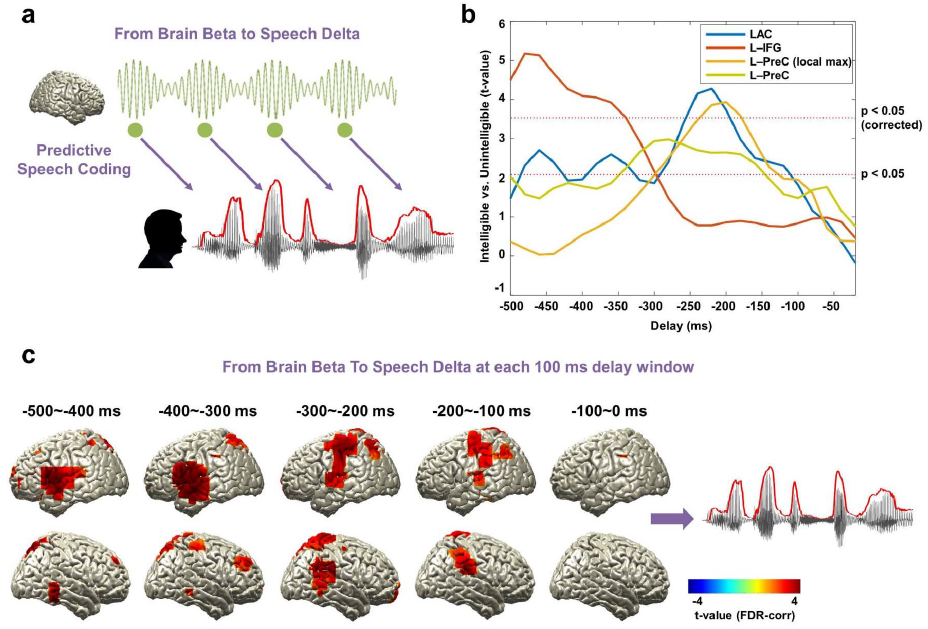
Beta rhythms and the prediction of upcoming speech. (A) A schematic figure for directed causal analysis: TE from brain beta power to speech delta phase (negative delays between speech-brain). TE computation was performed for each condition (intelligible-forward played and unintelligible-backward played) at each voxel from 20 ms to 500 ms in 20 ms steps and compared statistically between conditions. (b) TE values are averaged within each ROI from the AAL atlas and compared statistically between conditions. T-values are shown in each ROI: left primary auditory cortex (Heschl gyrus) (blue line), left IFG (orange line), and left precentral gyrus (both at local maximum coordinate (yellow line) and whole precentral gyrus ROI (light green line)). Statistical significance was shown with two red lines depicting t-values by paired t-test (upper red line: t_21_ = 3.53, p < 0.05, corrected; bottom red line: t_21_ = 2.08, p < 0.05, uncorrected). Left-lateralised predictions by beta power were observed (see Supplementary Figure 2). (c) Statistical contrast maps of averaged across 100 ms windows show that intelligibility-dependant prediction first engages left inferior frontal gyrus about 300-500 ms prior to the speech followed by left precentral gyrus and left primary auditory area 200–300 ms prior to the speech (p < 0.05, FDR-corrected).

Figure 4B shows over a range of delays for each ROI to what extent the prediction of forthcoming speech depends on speech intelligibility.

Beta power in left inferior frontal gyrus shows strong intelligibility-dependant prediction about 340-500 ms prior to speech (Figure 4B orange line). Left primary auditory cortex and left precentral gyrus show a similar significant effect but at a shorter delay peaking just before −200 ms (Figure 4B, blue and yellow lines). For the precentral gyrus, the ROI map from the AAL atlas is rather big, so we extracted the time series for the voxel with the local maximum in this area (yellow line, but also see the similar pattern for whole precentral gyrus ROI in the AAL atlas; light green line). Interestingly, this intelligibility-dependant prediction in these ROIs was left-lateralized (see Figure 4C and Supplementary Figure 2). Whole brain results averaged across 100 ms-long windows corroborated this pattern that first engages left inferior frontal gyrus about 300-500 ms prior to the speech followed by left precentral gyrus and left primary auditory area 200–300 ms prior to the speech (Figure 4C; p < 0.05, FDR-corrected).

### Relationship between temporal dynamics of speech entrainment and predictive speech coding

We next assessed how intelligibility-dependant prediction and the temporal dynamics of speech entrainment are related. We hypothesized that predictive control mechanisms interact with the rhythms in the brain driven by speech. We performed correlation analysis between TE values of intelligible speech at negative delays (preceding speech) and positive delays (following speech) using robust Spearman rank correlations across participants (Pernet, Wilcox, & Rousselet, 2012).

Motivated by the role of top-down beta activity particularly in the perception of sustained temporal aspect of speech (Pefkou, et al., 2017), we correlated predictive coding by beta activity (Figure 4) and speech entrainment (Figure 2) at various delays. We studied this mechanism first within early sensory area (auditory cortex) as well as higher order areas where we found strong top-down prediction.

Figure 5A shows this relationship in the left primary auditory cortex where predictive speech coding mechanism of beta power in the left auditory cortex ~250 ms prior to the forthcoming speech with low frequency delta phase information (corresponding to the plot for the left auditory cortex (blue line) at ~250 ms in the Figure 4B) is closely associated with low-frequency delta rhythm in the left auditory cortex driven by speech (corresponding to the plot for the left auditory cortex (pink line) at ~250 ms in the Figure 2B) (r = 0.46, p = 0.02). This suggests that individuals with stronger predictive coding by beta power in the left auditory cortex ~250 ms prior to the upcoming low frequency delta phase information in the speech signal are capable of stronger speech entrainment in the left auditory cortex by speech rhythm with low frequency delta phase at ~250 ms.

**Figure 5:**
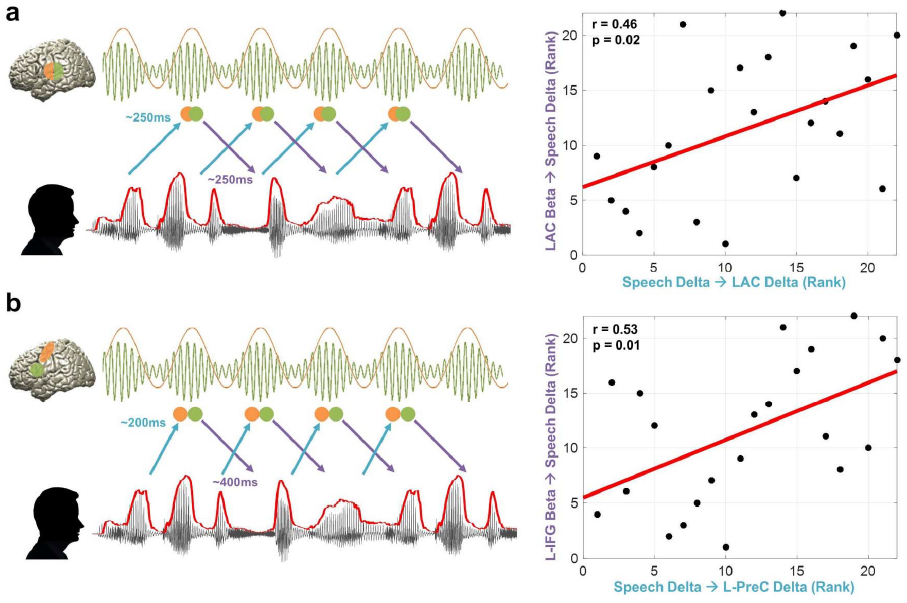
Relationship between temporal dynamics of speech entrainment and predictive speech coding. To assess how the temporal dynamics of speech entrainment (positive delays) and intelligibility-dependant top-down prediction (negative delays) are interacting, we performed correlation analysis using robust Spearman rank correlations across participants between the two mechanisms (but for top-down prediction, we used TE from brain beta to speech delta; Figure 4). We tested the relationship within in early sensory area. i.e. primary auditory cortex as well as higher order areas. (a) Delta phase in the left auditory cortex driven by speech (corresponding to the plot for the left auditory cortex (pink line) at ~250 ms in the Figure 2B) is associated with predictive speech coding mechanism of beta power in the left auditory cortex ~250 ms prior to the forthcoming speech with low frequency delta phase information (corresponding to the plot for the left auditory cortex (blue line) at ~250 ms in the Figure 4B) (r = 0.46, p = 0.02). (b) Delta phase in the left precentral gyrus modulated by the same frequency phase in the speech at ~200 ms (corresponding to the plot for the left precentral gyrus (pink line) at ~200 ms in the Figure 2D) is associated with predictive speech coding of beta power in the left inferior frontal gyrus ~400 ms prior to the forthcoming speech with low frequency delta phase information (corresponding to the plot for left inferior frontal gyrus (orange line) at ~400 ms in the Figure 4B) (r = 0.53, p = 0.01).

Another interesting aspect of this relationship emerged between the left inferior frontal cortex and left precentral gyrus. Figure 5B shows that predictive speech coding of beta power in the left inferior frontal gyrus ~400 ms prior to the forthcoming speech with low frequency delta phase information (corresponding to the plot for left inferior frontal gyrus (orange line) at ~400 ms in the Figure 4B) is associated with low frequency delta phase in the left precentral gyrus modulated by the same frequency phase in the speech at ~200 ms (corresponding to the plot for the left precentral gyrus (pink line) at ~200 ms in the Figure 2D) (r = 0.53, p = 0.01). This suggests that individuals with stronger modulation of predictive coding by beta power in the left inferior frontal gyrus ~400 ms prior to the low frequency delta phase information in the speech signal are capable of stronger speech entrainment in the left precentral gyrus by speech rhythm with low frequency delta phase at ~200 ms.

## Discussion

Here, we aimed to study the spatio-temporal characteristics of predictions during continuous speech recognition. We analysed transfer entropy (TE) at various (positive and negative) delays between the speech envelope and brain activity. At low frequencies (1-4 Hz) our results reveal a progression of predictive effects from left fronto-motor regions to right temporal regions. A different pattern emerged when we investigated TE between beta power in the brain and the phase of 1-4 Hz components in the speech envelope. We first see an engagement of left inferior frontal gyrus about 300-500 ms prior to speech followed by left precentral gyrus and left primary auditory areas. Our results suggest a progression of prediction processes from higher-order to early sensory areas.

First, it is important to carefully consider what aspects of predictions are captured using our approach. Transfer entropy (TE) is an information theoretic measure that quantifies directed statistical dependencies between time series. Specifically, TE from signal X to signal Y quantifies to what extent knowledge of X reduces uncertainty in predicting the future of Y over and above what could be predicted from knowledge of the past of Y alone. TE is conceptually similar to Granger causality as it infers causal relationships from time-lagged predictability. Here, when analysing prediction effects, we quantify to what extent the past of brain activity in a certain brain area improves prediction of the future of the speech envelope (over and above what could be predicted from the past of the speech envelope alone). Our main conclusions are then based on the statistical contrast between intelligible (forward played) speech and unintelligible (backward played) speech. This is important for two reasons. First, the power spectrum of the speech envelope is the same for both conditions. Therefore, this statistical contrast controls (to some extent) for the low-level rhythmicity in speech and amplifies sensitivity of the analysis to intelligibility. However, we acknowledge that backward played speech is not a perfect control condition due to differences in the finer temporal structure and attention to the stimulus. Second, similar to Granger causality, the computation of TE on bandpass-filtered data is not without problems (Florin, Gross, Pfeifer, Fink, & Timmermann, 2010; Weber, Florin, von Papen, & Timmermann, 2017). Statistically contrasting two conditions will counteract these problems. In addition, we would like to note that our results are very similar when using delayed mutual information (data not shown), a measure that is less sensitive to the effects of filtering. In summary, our approach is expected to be mostly sensitive to intelligibility-related components in speech.

Still, from our study it is difficult to exactly specify the structure in speech that is the target of the prediction processes presented here. This is in contrast to the many studies that demonstrate that a semantic violation at a certain point in a sentence gives rise to the well-known N400 evoked response (e.g. Kutas and Federmeier (2011)). While being less controlled, our approach benefits from ecological validity and more directly taps into prediction processes that operate during natural speech processing.

Brain oscillations in this frequency range follow the same frequency in the speech envelope robustly across all delays between 20-500 ms. The pattern is stronger in the right hemisphere (higher t-values in the Supplementary Figure 1). In our previous study we showed more right-lateralised speech-brain coupling at a fixed delay using mutual information (Figure 2A in Gross, Hoogenboom, et al. (2013)). This mechanism seems to extend to the directed TE measure used here up to ~400 ms delays. This supports the asymmetric sampling in time (AST) model (Poeppel, 2003) that posits a right hemisphere preference for longer temporal integration window (~150–250 ms). The sustained pattern suggests brain responses modulated by speech envelope are critical to continuous intelligible speech perception.

Directed coupling between speech envelope and brain oscillations across negative delays suggests a predictive coding mechanism where sensory processing is modulated in top-down manner. We investigated this mechanism from low-frequency delta (1-4 Hz) phase in the brain to low-frequency delta phase in the speech envelope and from beta power (14-18 Hz) in the brain to low-frequency delta (1-4 Hz) phase in the speech envelope. Our study reveals robust prediction processes at low-frequencies (1-4 Hz). This is the frequency that represents intonation and prosody (Ghitza, 2011; Giraud and Poeppel, 2012) but also overlaps at the upper end with the mean syllable rate (Ding et al., 2017). It shows an interesting temporal progression from left inferior fronto-motor areas to right auditory-temporal areas. This progression of prediction from higher order areas in the left hemisphere (200–300 ms prior to speech) to the early sensory areas in the right hemisphere suggests that the brain first generates prediction of upcoming sensory input by top-down contextual knowledge that is later used for optimised stimulus encoding in early sensory areas. Indeed, top-down modulations of delta phase (such as temporal expectation or selection of an attended stimulus stream) has been shown to increase sensitivity to external inputs in the auditory (Lakatos, Karmos, Mehta, Ulbert, & Schroeder, 2008) and visual domain (Cravo, Rohenkohl, Wyart, & Nobre, 2013). Similarly, we find that beta power in the left frontal cortex and sensorimotor areas reflects prediction of upcoming speech relatively early (200–500 ms prior to speech) and is left-lateralised (Supplementary Figure 2). Active inference by motor systems regarding predictive coding has been studied recently and beta oscillation has been suggested to be working together with low-frequency activity in top-down modulation of ongoing activity during predictive coding (Arnal and Giraud, 2012). Recently an elegant study employing an auditory attention task has shown that interdependent delta and beta activity from left sensorimotor cortex encodes temporal prediction and this is directed towards auditory areas (Morillon and Baillet, 2017). This is consistent with our finding that both delta phase and beta power in the left frontal and sensorimotor engages in the prediction of forthcoming speech from relatively early stage. The temporal progression from inferior frontal to motor areas seems to suggest a hierarchical organisation of prediction processes that warrant further investigation.

During continuous speech perception, brain oscillations entrained by speech (positive delays) and predicting speech (negative delays) are expected to interact in time. In other words, there is a continuous recurrent interaction between stimulus-driven bottom-up processing and top-down prediction processing during continuous speech perception. We focused our analysis on the interaction between top-down predictive coding, i.e., TE from beta power in the brain to delta phase in the speech (Figure 4, negative delays) in the auditory cortex and higher order areas, i.e., fronto-motor areas and speech entrainment, i.e., TE from delta phase in the speech to delta phase in the brain (Figure 2, positive delays). In the left auditory cortex (Figure 5A), subjects with better top-down prediction by beta power at ~250 ms (~ peak of LAC in Figure 4B; Supplementary Figure 2B) show better entrainment by speech delta phase at ~250 ms (~crossing point between the LAC and RAC in Figure 2B). This result indicates that although low-frequency brain oscillations following speech envelope seems stronger in the right hemisphere than the left hemisphere (higher t-value for RAC at early delays in Figure 2B; Supplementary Figure 1), the interaction between both (top-down prediction and bottom-up speech entrainment) is modulated by the left auditory cortex (Park, et al., 2015). In the higher order areas (Figure 5B), speech-driven bottom-up information flow to the left motor cortex in the delta phase at delays ~200 ms is strongly associated with top-down predictive information flow from beta power in the left IFG to speech delta phase at ~400 ms prior to the speech. This indicates that subjects with better ability to control top-down predictive speech coding ~400 ms prior to the upcoming speech by beta power in the left IFG (orange line in Figure 4B; Supplementary Figure 2C) are also better bottom-up entrained by speech delta phase at ~200 ms in the left motor cortex (pink line in Figure 2D).

In summary, our results indicate that predictive processes during continuous speech processing involve fronto-motor areas, operate in at least two frequency channels (delta and beta), follow an organised temporal progression from higher-order areas to early sensory areas and recurrently interact with reactive processes. Further research is needed to decode the exact nature of these predictions, identify the contributions of individual areas and elucidate the mutual dependencies between processes that precede and follow speech.

## Acknowledgments

JG is supported by the Wellcome Trust (098433) and GT is supported by the Wellcome Trust (098434). The funders had no role in study design, data collection and analysis, decision to publish, or preparation of the manuscript.

## Supplemental Materials

**Supplemental Figure 1:**
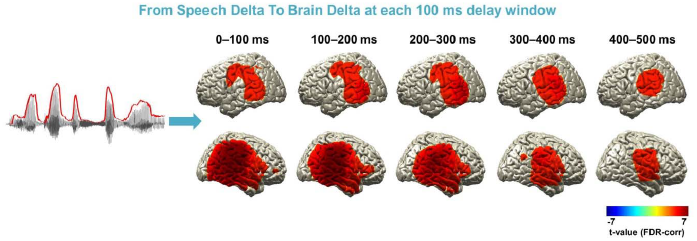
(Related to Figure 2). Entrained brain signals following speech at whole brain at each 100 ms window. TE computation was performed for each condition (intelligible-forward played and unintelligible-backward played) at each voxel from 20 ms to 500 ms with a 20-ms step. To characterise the spatio-temporal pattern across delays at whole brain level, we averaged TE-maps in 100 ms windows and computed again the statistical contrast of intelligible compared to unintelligible speech (only t-values > 4.7, p < 0.0001, FDR-corrected are plotted). Delta phase information in the brain following the same frequency information in the speech envelope is robust across all delays and the pattern is stronger in the right hemisphere (higher t-values).

**Supplemental Figure 2:**
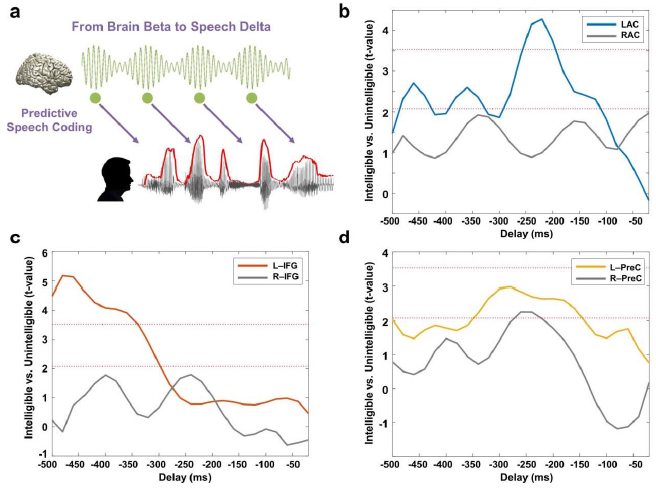
Supplemental Figure 2 (related to Figure 4). Beta rhythms involving the prediction of upcoming speech are left-lateralised. Here we show the same plots as in the Figure 4B, but separately for each ROI with the homologous ROI in the right hemisphere; (a) Primary auditory cortex (Heschl gyrus), (b) Inferior frontal gyrus – opercular part, (c) Precentral gyrus. Top-down prediction by beta power in the brain to speech delta phase is left-lateralised (statistics by paired t-test; upper red line: t_21_ = 3.53, p < 0.05, corrected; bottom red line: t_21_ = 2.08, p < 0.05, uncorrected). Our results indicate three different mechanisms in terms of hemispheric asymmetry. 1) Speech-driven entrainment by delta phase is shown bilaterally (Figure 2; Supplementary Figure 1). 2) However, top-down prediction by the same delta phase shows progression from left inferior fronto-motor areas (200–300 ms prior to speech) to right auditory-temporal areas (0–200 ms prior to speech). 3) Top-down prediction by beta power in the fronto-motor areas is left-lateralised from early stage (200–500 ms prior to speech).

## References

Arnal, L. H., & Giraud, A. L. (2012). Cortical oscillations and sensory predictions. [Review]. Trends Cogn Sci, 16(7), pp. 390–398. doi:10.1016/j.tics.2012.05.003 Retrieved from http://www.ncbi.nlm.nih.gov/pubmed/22682813

Bastos, A. M., Usrey, W. M., Adams, R. A., Mangun, G. R., Fries, P., & Friston, K. J. (2012). Canonical microcircuits for predictive coding. Neuron, 76(4), pp. 695–711. doi:10.1016/j.neuron.2012.10.038 Retrieved from http://www.ncbi.nlm.nih.gov/pubmed/23177956

Besl, P. J., & McKay, N. D. (1992). A method for registration of 3-D shapes. IEEE T Pattern Anal, pp.239–256.

Brainard, D. H. (1997). The Psychophysics Toolbox. Spat Vis, 10(4), pp. 433–436. Retrieved from http://www.ncbi.nlm.nih.gov/pubmed/9176952

Chandrasekaran, C., Trubanova, A., Stillittano, S., Caplier, A., & Ghazanfar, A. A. (2009). The natural statistics of audiovisual speech. PLoS Comput Biol, 5(7), p e1000436. doi:10.1371/journal.pcbi.1000436 Retrieved from http://www.ncbi.nlm.nih.gov/pubmed/19609344

Cravo, A. M., Rohenkohl, G., Wyart, V., & Nobre, A. C. (2013). Temporal expectation enhances contrast sensitivity by phase entrainment of low-frequency oscillations in visual cortex. J Neurosci, 33(9), pp. 4002–4010. doi:10.1523/JNEUROSCI.4675-12.2013 Retrieved from https://www.ncbi.nlm.nih.gov/pubmed/23447609

Ding, N., Melloni, L., Zhang, H., Tian, X., & Poeppel, D. (2016). Cortical tracking of hierarchical linguistic structures in connected speech. Nat Neurosci, 19(1), pp. 158–164. doi:10.1038/nn.4186 Retrieved from https://www.ncbi.nlm.nih.gov/pubmed/26642090

Ding, N., Patel, A. D., Chen, L., Butler, H., Luo, C., & Poeppel, D. (2017). Temporal modulations in speech and music. Neurosci Biobehav Rev, 81(Pt B), pp. 181–187. doi:10.1016/j.neubiorev.2017.02.011 Retrieved from https://www.ncbi.nlm.nih.gov/pubmed/28212857

Florin, E., Gross, J., Pfeifer, J., Fink, G. R., & Timmermann, L. (2010). The effect of filtering on Granger causality based multivariate causality measures. Neuroimage, 50(2), pp. 577–588. doi:10.1016/j.neuroimage.2009.12.050 Retrieved from https://www.ncbi.nlm.nih.gov/pubmed/20026279

Fontolan, L., Morillon, B., Liegeois-Chauvel, C., & Giraud, A. L. (2014). The contribution of frequency-specific activity to hierarchical information processing in the human auditory cortex. Nat Commun, 5, p 4694. doi:10.1038/ncomms5694 Retrieved from http://www.ncbi.nlm.nih.gov/pubmed/25178489

Friston, K. J., & Frith, C. D. (2015). Active inference, communication and hermeneutics. Cortex, 68, pp. 129–143. doi:10.1016/j.cortex.2015.03.025 Retrieved from https://www.ncbi.nlm.nih.gov/pubmed/25957007

Ghitza, O. (2011). Linking speech perception and neurophysiology: speech decoding guided by cascaded oscillators locked to the input rhythm. Front Psychol, 2, p 130. doi:10.3389/fpsyg.2011.00130 Retrieved from http://www.ncbi.nlm.nih.gov/pubmed/21743809

Giordano, B. L., Ince, R. A. A., Gross, J., Schyns, P. G., Panzeri, S., & Kayser, C. (2017). Contributions of local speech encoding and functional connectivity to audio-visual speech perception. Elife, 6doi:10.7554/eLife.24763 Retrieved from https://www.ncbi.nlm.nih.gov/pubmed/28590903

Giraud, A. L., & Poeppel, D. (2012). Cortical oscillations and speech processing: emerging computational principles and operations. [Research Support, N.I.H., Extramural Research Support, Non-U.S. Gov’t]. Nat Neurosci, 15(4), pp. 511–517. doi:10.1038/nn.3063 Retrieved from http://www.ncbi.nlm.nih.gov/pubmed/22426255

Gross, J., Baillet, S., Barnes, G. R., Henson, R. N., Hillebrand, A., Jensen, O., … Schoffelen, J. M. (2013). Good practice for conducting and reporting MEG research. [Research Support, Non-U.S. Gov’t]. Neuroimage, 65, pp. 349–363. doi:10.1016/j.neuroimage.2012.10.001 Retrieved from http://www.ncbi.nlm.nih.gov/pubmed/23046981

Gross, J., Hoogenboom, N., Thut, G., Schyns, P., Panzeri, S., Belin, P., & Garrod, S. (2013). Speech rhythms and multiplexed oscillatory sensory coding in the human brain. PLoS Biol, 11(12), p e1001752. doi:10.1371/journal.pbio.1001752 Retrieved from http://www.ncbi.nlm.nih.gov/pubmed/24391472

Ince, R. A., Mazzoni, A., Bartels, A., Logothetis, N. K., & Panzeri, S. (2012). A novel test to determine the significance of neural selectivity to single and multiple potentially correlated stimulus features. J Neurosci Methods, 210(1), pp. 49–65. doi:10.1016/j.jneumeth.2011.11.013 Retrieved from https://www.ncbi.nlm.nih.gov/pubmed/22142889

Ince, R. A., Schultz, S. R., & Panzeri, S. (2014). Estimating Information-Theoretic Quantities. In D. Jaeger & R. Jung (Eds.), Encyclopedia of Computational Neuroscience.

Kutas, M., & Federmeier, K. D. (2011). Thirty years and counting: finding meaning in the N400 component of the event-related brain potential (ERP). Annu Rev Psychol, 62, pp. 621–647. doi:10.1146/annurev.psych.093008.131123 Retrieved from https://www.ncbi.nlm.nih.gov/pubmed/20809790

Lakatos, P., Karmos, G., Mehta, A. D., Ulbert, I., & Schroeder, C. E. (2008). Entrainment of neuronal oscillations as a mechanism of attentional selection. Science, 320(5872), pp. 110–113. doi:10.1126/science.1154735 Retrieved from http://www.ncbi.nlm.nih.gov/pubmed/18388295

Levinson, S. C. (2016). Turn-taking in Human Communication--Origins and Implications for Language Processing. Trends Cogn Sci, 20(1), pp. 6–14. doi:10.1016/j.tics.2015.10.010 Retrieved from https://www.ncbi.nlm.nih.gov/pubmed/26651245

Massey, J. (1990). Causality, feedback and directed information. In: Proc. Int. Symp. Information Theory Application (ISITA 1990), pp. 303–305.

Morillon, B., & Baillet, S. (2017). Motor origin of temporal predictions in auditory attention. Proc Natl Acad Sci U S A, 114(42), pp. E8913–E8921. doi:10.1073/pnas.1705373114 Retrieved from https://www.ncbi.nlm.nih.gov/pubmed/28973923

Nolte, G. (2003). The magnetic lead field theorem in the quasi-static approximation and its use for magnetoencephalography forward calculation in realistic volume conductors. Phys Med Biol, 48(22), pp. 3637–3652. Retrieved from http://www.ncbi.nlm.nih.gov/pubmed/14680264

Norris, D., McQueen, J. M., & Cutler, A. (2016). Prediction, Bayesian inference and feedback in speech recognition. Lang Cogn Neurosci, 31(1), pp. 4–18. doi:10.1080/23273798.2015.1081703 Retrieved from https://www.ncbi.nlm.nih.gov/pubmed/26740960

Oostenveld, R., Fries, P., Maris, E., & Schoffelen, J. M. (2011). FieldTrip: Open source software for advanced analysis of MEG, EEG, and invasive electrophysiological data. [Research Support, Non-U.S. Gov’t]. Comput Intell Neurosci, 2011, p 156869. doi:10.1155/2011/156869 Retrieved from http://www.ncbi.nlm.nih.gov/pubmed/21253357

Panzeri, S., Senatore, R., Montemurro, M. A., & Petersen, R. S. (2007). Correcting for the sampling bias problem in spike train information measures. J Neurophysiol, 98(3), pp. 1064–1072. doi:10.1152/jn.00559.2007 Retrieved from https://www.ncbi.nlm.nih.gov/pubmed/17615128

Park, H., Ince, R. A., Schyns, P. G., Thut, G., & Gross, J. (2015). Frontal Top-Down Signals Increase Coupling of Auditory Low-Frequency Oscillations to Continuous Speech in Human Listeners. Curr Biol, 25(12), pp. 1649–1653. doi:10.1016/j.cub.2015.04.049 Retrieved from http://www.ncbi.nlm.nih.gov/pubmed/26028433

Park, H., Kayser, C., Thut, G., & Gross, J. (2016). Lip movements entrain the observers’ low-frequency brain oscillations to facilitate speech intelligibility. Elife, 5doi:10.7554/eLife.14521 Retrieved from https://www.ncbi.nlm.nih.gov/pubmed/27146891

Pefkou, M., Arnal, L. H., Fontolan, L., & Giraud, A. L. (2017). theta-Band and beta-Band Neural Activity Reflects Independent Syllable Tracking and Comprehension of Time-Compressed Speech. J Neurosci, 37(33), pp. 7930–7938. doi:10.1523/JNEUROSCI.2882-16.2017 Retrieved from https://www.ncbi.nlm.nih.gov/pubmed/28729443

Pernet, C. R., Wilcox, R., & Rousselet, G. A. (2012). Robust correlation analyses: false positive and power validation using a new open source matlab toolbox. Front Psychol, 3, p 606. doi:10.3389/fpsyg.2012.00606 Retrieved from https://www.ncbi.nlm.nih.gov/pubmed/23335907

Pickering, M. J., & Garrod, S. (2007). Do people use language production to make predictions during comprehension? Trends Cogn Sci, 11(3), pp. 105–110. doi:10.1016/j.tics.2006.12.002 Retrieved from http://www.ncbi.nlm.nih.gov/pubmed/17254833

Pickering, M. J., & Garrod, S. (2013). An integrated theory of language production and comprehension. Behav Brain Sci, 36(4), pp. 329–347. doi:10.1017/S0140525X12001495 Retrieved from http://www.ncbi.nlm.nih.gov/pubmed/23789620

Poeppel, D. (2003). The analysis of speech in different temporal integration windows: cerebral lateralization as ‘asymmetric sampling in time’. Speech Communication, 41, pp. 245–255.

Schreiber, T. (2000). Measuring information transfer. Phys Rev Lett, 85(2), pp. 461–464. doi:10.1103/PhysRevLett.85.461 Retrieved from https://www.ncbi.nlm.nih.gov/pubmed/10991308

Tzourio-Mazoyer, N., Landeau, B., Papathanassiou, D., Crivello, F., Etard, O., Delcroix, N., … Joliot, M. (2002). Automated anatomical labeling of activations in SPM using a macroscopic anatomical parcellation of the MNI MRI single-subject brain. Neuroimage, 15(1), pp. 273–289. doi:10.1006/nimg.2001.0978 Retrieved from https://www.ncbi.nlm.nih.gov/pubmed/11771995

Weber, I., Florin, E., von Papen, M., & Timmermann, L. (2017). The influence of filtering and downsampling on the estimation of transfer entropy. PLoS One, 12(11), p e0188210. doi:10.1371/journal.pone.0188210 Retrieved from https://www.ncbi.nlm.nih.gov/pubmed/29149201

Zion Golumbic, E., Cogan, G. B., Schroeder, C. E., & Poeppel, D. (2013). Visual input enhances selective speech envelope tracking in auditory cortex at a cocktail party. J Neurosci, 33(4), pp. 1417–1426. doi:10.1523/JNEUROSCI.3675-12.2013 Retrieved from http://www.ncbi.nlm.nih.gov/pubmed/23345218

